# Spontaneous Necker-Cube Reversals are not that Spontaneous

**DOI:** 10.1101/2023.03.06.531257

**Authors:** Mareike Wilson, Lukas Hecker, Ellen Joos, Ad Aertsen, Ludger Tebartz van Elst, Jürgen Kornmeier

## Abstract

During observation of the ambiguous Necker cube, our perception suddenly reverses between two about equally possible 3D interpretations. During passive observation, perceptual reversals seem to be sudden and spontaneous. A number of theoretical approaches postulate destabilization of neural representations as a precondition for spontaneous reversals of ambiguous figures. In the current study, we focused on possible EEG correlates of perceptual destabilization, that may allow to predict an upcoming perceptual reversal.

We presented ambiguous Necker cube stimuli in an onset-paradigm and investigated the neural processes underlying endogenous reversals as compared to perceptual stability across two consecutive stimulus presentations. In a separate experimental condition, disambiguated cube variants were alternated randomly, to exogenously induce perceptual reversals. We compared the EEG immediately before and during endogenous Necker cube reversals with corresponding time windows during exogenously induced perceptual reversals of disambiguated cube variants.

For the ambiguous Necker cube stimuli, we found the earliest differences in the EEG between reversal trials and stability trials already one second before a reversal occurred, at bilateral parietal electrodes. The traces remained similar until approximately 1100 ms before a perceived reversal, became maximally different at around 890 ms (p=7.59*10^−6^, Cohen’s d=1.35) and remained different until shortly before offset of the stimulus preceding the reversal. No such patterns were found in the case of disambiguated cube variants.

The identified EEG effects may reflect destabilized states of neural representations, related to destabilized perceptual states preceding a perceptual reversal. They further indicate that spontaneous Necker cube reversals are most probably not as spontaneous as generally thought. Rather, the destabilization may occur over a longer time scale, at least one second before a reversal event.

## 1 Introduction

During observation of an ambiguous figure, like the famous Necker cube (Necker, 1832), our perception becomes unstable and alternates between two, or even more, possible interpretations despite unchanged visual input. Observers can volitionally control their percept to some degree and both increase and decrease their reversal rate (the number of perceptual reversals over time), however, they cannot prevent the reversals entirely (e.g., Kornmeier et al., 2009, 2019; Strüber & Stadler, 1999; van Ee et al., 2005). Multistable perception can also be induced via binocular rivalry, i.e. if the two eyes simultaneously receive conflicting information. Similar to the situation with classical ambiguous figures, perception alternates between the two percepts corresponding to the two eyes’ input (e.g. Blake, 2001; O’Shea et al., 2013). Multistable perception phenomena have also been reported in other modalities, like audition (e.g., Einhäuser et al., 2020; Pressnitzer & Hupe, 2006) and touch (e.g., Carter et al., 2008; Conrad et al., 2012; Darki & Rankin, 2020; Hense et al., 2019; Liaci et al., 2016).

The phenomenon of multistable perception has fascinated researchers from different disciplines for more than 200 years (J. Brascamp et al., 2018). Multistable perception is vividly discussed in the contexts of perception and consciousness (e.g., Crick & Koch, 1998; Blake & Logothetis, 2002; Long & Toppino, 2004; Atmanspacher et al., 2008; J. Brascamp et al., 2018; Devia et al., 2022; Giles et al., 2016), particularly because a separation of sensory processing (which can be kept constant over time) and perceptual processing (which alternates repeatedly) (e.g., Bartels, 2021; Blake et al., 2014; Crick & Koch, 1998; Intaite et al., 2010; Leopold & Logothetis, 1999; Tong et al., 1998) appears to be possible. However, despite a large number of experimental studies, the neural mechanisms underlying spontaneous perceptual reversals are so far poorly understood. Problems are both theoretical and practical in nature. Very often, it is unclear what the processes are that (1) necessarily precede a spontaneous perceptual reversal (e.g. the weakening or destabilization of a neural representation) and causally lead to it, and (2) those of the reversal process as such, i.e. the change in dominance (and access to consciousness) of one neural representation by the alternative one in a certain time window, and finally (3) the subsequent processes, related to becoming aware of the reversal event and communicating it with the environment within an experimental paradigm (J. Brascamp et al., 2018; Devia et al., 2022). The EEG provides the necessary temporal resolution in the range of milliseconds to differ between these processing steps. However, the precise temporal reference of the reversal event is necessary for the differentiation between the above-described processing steps but at the same time difficult to access due to its endogenous nature.

A number of EEG- and fMRI-studies used participants’ key presses as a time reference for the reversal event (e.g., Basar-Eroglu et al., 1996; Knapen et al., 2011; Lumer et al., 1998; Sterzer & Kleinschmidt, 2007). This method, however, comes with the problem of intra-individual reaction time variability in the range of ± 100 ms (Kornmeier & Bach, 2004, 2012; Strüber & Herrmann, 2002), which is too large to exploit the high temporal resolution of the EEG. An alternative approach used ambiguous motion stimuli with specific stimulus features or presentation modes that allow to narrow down a possible time window of the reversal event (e.g, Pastukhov, 2016; Pastukhov & Braun, 2013; Weilnhammer et al., 2017, 2021). This elegant stimulus design, however, is restricted to a certain class of motion stimuli. De Jong et al. recently developed a clever method using within-participant intracranial electrophysiological measures to temporally resolve reversals of binocular rivalry stimuli (de Jong et al., 2020).

Other studies referred to the change in pupil diameter during perceptual reversals (J. W. Brascamp et al., 2021; Einhäuser et al., 2008; Hupe et al., 2009; Kloosterman et al., 2015) or used the horizontal eye fixation positions in a free observation paradigm (Polgári et al., 2020). These innovative approaches allowed the implementation of “no report paradigms” and helped to separate task-related and perception-related processing steps (e.g., J. Brascamp et al., 2018; J. W. Brascamp et al., 2021; Devia et al., 2022). However, the specific role of the pupil response in the reversal process is discussed controversially (Duman et al., 2022). It is particularly unclear, whether the pupil response or a certain change in horizontal eye position precedes or follows the reversal event and how much the latency between the reversal event and the pupil response/eye position change varies across repetitions.

In the present study, we used the so-called “Onset Paradigm” to estimate the timing of the reversal event. The basic idea goes back to the seminal work of Orbach et al. (1963, 1966). They presented the Necker cube discontinuously and found a huge variability of the reversal rate (number of reversals per minute) as a function of the inter-stimulus interval (ISI), i.e. the duration of a blank screen between successive Necker cube presentations. The reversal rate continuously increased from continuous presentations to ISIs of about 400 milliseconds (ms) and decreased again for longer ISIs, up to a complete freezing of the percept for ISIs of about 2 seconds (Kornmeier et al., 2007; Leopold et al., 2002; Maier et al., 2003; Pastukhov & Braun, 2008; Pearson & Brascamp, 2008; Zaretskaya et al., 2010). O’Donnell (1988) were the first to present Necker cubes discontinuously while measuring EEG. They found a P300 event-related potential (ERP) component correlating with a perceptual reversal. Kornmeier et al. (2004) adopted this idea and developed the “Onset Paradigm”: Participants compared their percepts of subsequently presented Necker cubes and indicated in separate experimental conditions either perceptual reversals (reversal condition) or perceptual stability (stability conditions) from the previously observed stimulus to the current stimulus by a key press in the subsequent ISI. The idea of this paradigm was, that the discontinuous presentation mode synchronizes the reversal event with stimulus onset, allowing the latter as a time reference for the former. Kornmeier et al. calculated ERPs and subtracted stability trials from reversal trials in order to eliminate the ERP signatures related to stimulus onset, decision making and motor preparation.

In a separate experiment with the same conditions, Kornmeier and colleagues replaced the ambiguous Necker cube with disambiguated cube versions, corresponding to the two perceptual interpretations of the ambiguous Necker cube stimulus. Their Onset Paradigm increased the temporal resolution of the reversal event strongly and revealed two highly similar chains of ERP components for endogenous perceptual reversals (Experiment 1) and exogenously induced reversals (Experiment 2) with two exceptions. The ERP chain related to the endogenous reversals started with a small occipital positivity (“Reversal Positivity”) at 160 ms after stimulus onset. This Reversal Positivity was not observed with exogenously induced reversals. It may indicate an early visual “detection” of a maximally unstable perceptual state in the case of an endogenous reversal, as though the early perceptual processing units are not able to handle the sensory information in the first feed-forward loop of sensory information flow (e.g., Kornmeier et al., 2011). The exogenously induced reversal (realized by the computer program) is void of this problem, which may explain the absence of the Reversal Positivity in this case. Furthermore, the subsequent three ERP components related to endogenous reversals were delayed by about 40 to 60 ms compared to those from the exogenously induced reversals. A comparable processing delay was observed in the reaction time data in a separate experiment and may reflect the necessary time to solve the sensory ambiguity problem.

Since its introduction, the Onset Paradigm has been used by several labs around the world and the findings, reported above, have mainly been confirmed (Abdallah & Brooks, 2020; Atmanspacher et al., 2008; Britz et al., 2010; Ehm et al., 2011; Intaite et al., 2010, 2013; Kornmeier et al., 2007, 2011, 2019; Kornmeier & Bach, 2004, 2006, 2014; Pitts et al., 2007, 2008, 2010; Sandberg et al., 2014; Yokota et al., 2014). For reviews see Kornmeier & Bach (2012) and Pitts & Britz (2011).

For the interpretation of their ERP findings, Kornmeier et al. postulated two separate processes (see Fig. 2 for a schematic graphical representation): After a perceptual interpretation has been established, its neural representation starts to slowly destabilize until a state of maximal neural instability has been reached. Assuming that the brain has evolved to keep unstable states as short as possible, this instability becomes resolved by a fast restabilization process of 40 – 60 ms, as indicated by the ERP results, leading to the reversed perceptual interpretation (Kornmeier & Bach, 2006).

These previous EEG studies indicated that this restabilization is mainly executed in lower visual areas. However, it is so far unclear, which brain areas are affected during the postulated longer-lasting destabilization process and at which time points between two perceptual reversals we can find EEG correlates of a destabilized perceptual state. Hence, the aim of the present study is to identify EEG signatures that reflect states of gradual destabilization preceding a reversal event. Such signatures would allow us to better understand the processes underlying perceptual / neural destabilization. We focused on ERP differences between stable and unstable perceptual states in a time window before a reversal.

## 2 Methods

### 2.1 Participants

EEG data from 21 participants (mean age=25.0 years, standard deviation (SD)=3.27 years; 11 female) was collected and analyzed. All participants had normal or corrected-to-normal visual acuity as measured with the Freiburg Visual Acuity and Contrast Test (FrACT, Bach, 2018) and gave their informed written consent. The study was approved by the ethics committee of the University of Freiburg, Germany.

### 2.2 Stimuli

As visual stimuli we used the ambiguous Necker lattice (a combination of nine Necker cubes, Necker, 1832) as first described in Kornmeier and Bach (Kornmeier et al., 2001; Kornmeier & Bach, 2004) (Figure 1) and two disambiguated lattice variants corresponding to the two perceptual interpretations of the ambiguous lattice, including depth cues, like shading, central projection, and aerial perspective (Fig. 1). The lattices were white (40cd/m2) with a dark background (0.01 cd/m^2^). This was the luminance for both lattice variants, the disambiguated variant’s luminance was calculated by averaging the luminance across the four corners. The Necker lattices had a size of 7.5° × 7.5° degrees of visual angle. A fixation cross was presented in the center of the lattices (Joos et al., 2020).

**Figure 1.**
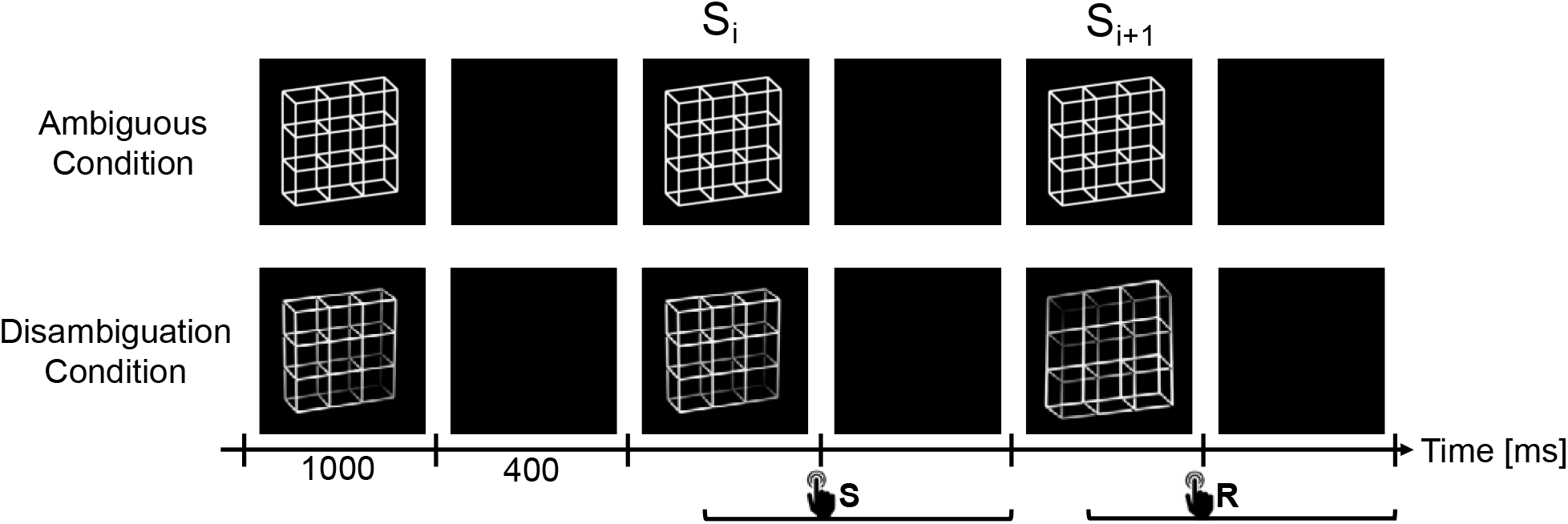
The experiment consisted of two experimental conditions, the Ambiguity Condition (top row) and Disambiguation Condition (bottom row). In both conditions, the lattice stimuli were presented for 1000 ms with ISIs of 400 ms in between. Participants had to compare successive stimuli and indicate via button press whether they perceived the current lattice orientation as reversed compared to the previous lattice (reversal trials, R) or as unchanged (stability trials, S). Participants were allowed to respond from stimulus onset until the end of the subsequent ISI. We focused our analysis on the stimulus time window before (S_i_) and after (S_i+1_) the reversed percept, including the ISI in between. Diagram adapted from Kornmeier and Bach (2012) and Joos et al. (2020).

**Figure 2.**
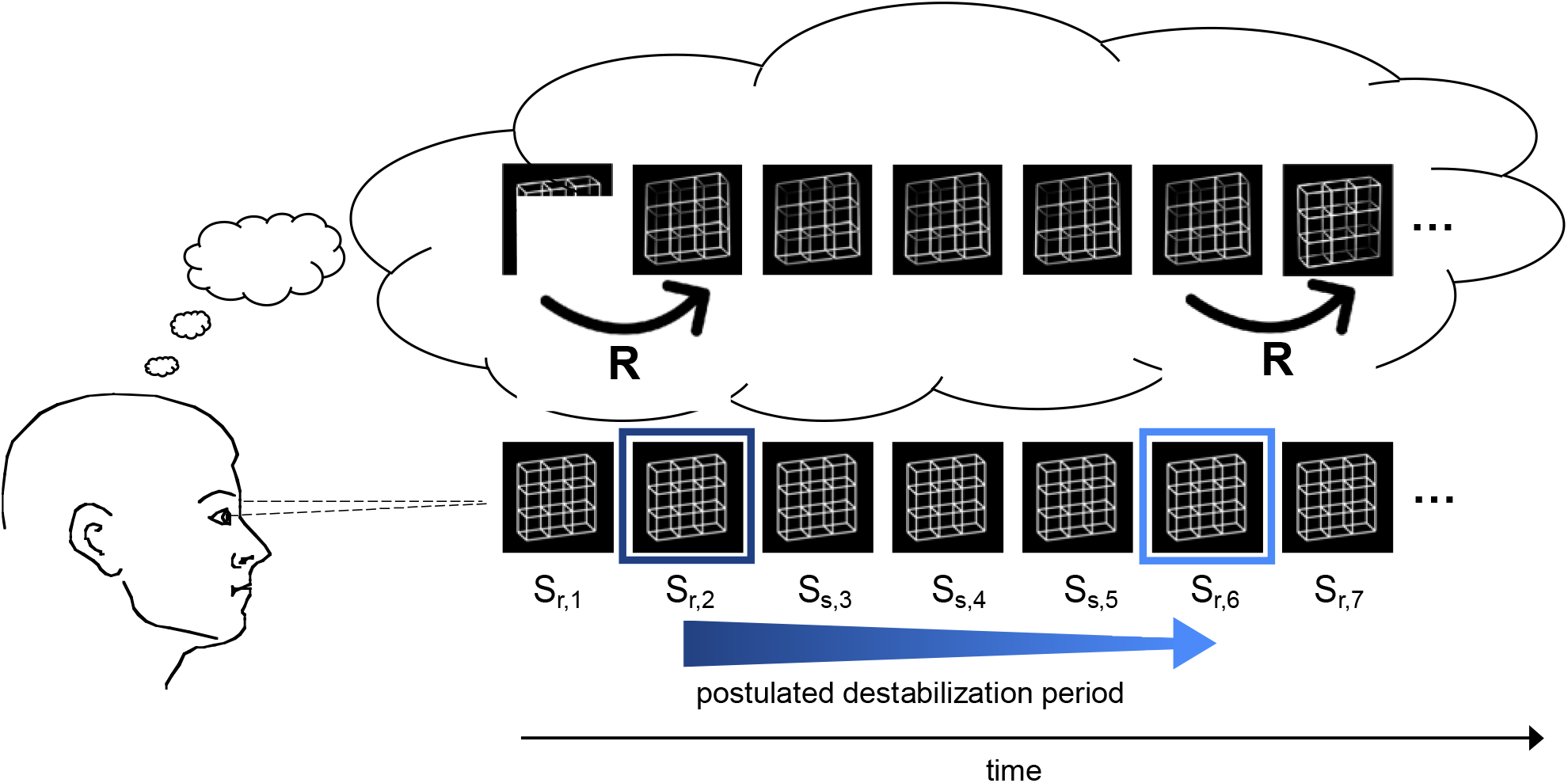
Schematic demonstration of the reversal dynamics. The participant observes an ambiguous Necker lattice discontinuously over a certain time interval and perceives it as facing in one orientation. The perceptual representation is postulated to destabilize over time (blue arrow) until the percept becomes so unstable (indicated by the light blue frame) that a perceptual reversal occurs. After the reversal, the perceptual representation is again temporally stable (dark blue frame). Bottom row: Necker lattice stimuli as ambiguous sensory input. Top row: perceptual interpretation of the observer. The instances R indicate perceptual reversals.

#### 2.2.1 Procedure

In two experimental conditions, either ambiguous or disambiguated variants of the Necker lattice were presented discontinuously (1000 ms presentation time) with blank screen ISIs (400 ms duration) between stimulus presentations. In a 1-back task, participants compared the perceived 3D orientation of the currently presented lattice stimulus (S_i+1_) with the previously presented stimulus (S_i_) and indicated perceptual reversals and perceptual stability (same percept across the two stimulus presentations) by different keys. We kept the stimulus presentation time short enough (1000 ms) to prevent additional reversals during stimulus observation. In the case of ambiguous lattice stimuli, perceptual reversals across stimuli were evoked endogenously by the participants perceptual system. In the case of the disambiguated lattice variants, the reversal was evoked by the computer program with a predefined reversal rate of 30%. The two experimental conditions were subdivided in experimental blocks of approximately 7 min duration, containing typically some 300 trials.

Before the experiment, participants completed a training session where they were presented with disambiguated Necker lattices. Identical to the experiment, participants needed to indicate with a button press if the percept stated the same or changed from S_i_ to S_i+1_. The training duration was approximately 5 minutes. If participants got less than 90% of the trials correct, the training was repeated.

##### 2.2.1.1 Reversal rates

We calculated the number of reversals per minute (reversal rate) by counting the number of reversals per experimental block and dividing the sum by the block duration. One block lasted for 7 minutes on average. This reversal rate was then averaged within participants across experimental blocks of the same condition. The focus of the present analysis was the identification of EEG signatures related to perceptual instability before a reversal compared to a time window of perceptual stability. For this purpose, it was necessary to have a minimum number of reversal trials per participant. We therefore excluded participants with less than 5 reversals per minute, resulting in a dataset that included 15 participants from the original 21.

### 2.3 EEG Recording

The EEG was recorded with 32 active silver/silver chloride electrodes using the extended 10-20 system for the electrode positions on the scalp (American Clinical Neurophysiology Society, 2006), using the BrainVision ActiCHamp amplifier. The data was digitized at a sampling rate of 1000 Hz and online bandpass filtered at 0.01-120Hz. Impedance was kept below 10 kΩ for all electrodes.

### 2.4 Preprocessing

All preprocessing was done using the MNE-Python package (version 0.23.0) (Gramfort et al., 2013) in Python 3.9.6. The raw data was offline band-pass filtered from 0.01 to 25 Hz and re-referenced to the average of the mastoid electrodes (TP9, TP10). The vertical electrooculogram (vEOG) electrodes were used to determine eye artifacts. Due to technical problems, one participant did not have a vEOG electrode, and therefore the Fp1 electrode was used for artifact detection. The vEOG electrode (or Fp1 electrode) was bandpass filtered from 1 to 10 Hz.

To detect eye movement artifacts an Independent Component Analysis (ICA) was conducted. A correlation was calculated between the vEOG electrode and the Independent Components (IC) to detect which components were most correlated to eye movements. If a component surpassed a predefined correlation threshold of r=0.8, this component was eliminated from the data.

For the remaining artifacts an artifact rejection threshold of ± 100µV (peak-to-peak amplitude) was defined. Data was baseline corrected, with the average amplitude in a time window between 60 ms before to 40 ms after stimulus onset of S_i+1_ as baseline. Trials were labelled as either reversal or stable depending on the participants’ responses. Responses were regarded as valid if given in a time from 150 ms after S_i+1_ onset to the end of the ISI following S_i+1_. If a response button was pressed more than once during this response time interval, the latest response was used. In cases of pressing wrong buttons or no responses the trial was ignored.

### 2.5 Data Analysis

For the data analysis we collected reversal trials S_r,i+1_, where the perceptual reversal took place from stimulus S_r,i_ to stimulus S_r,i+1._ We compared these reversal trials with stability trials S_s,i+1_, where perception stayed unchanged from S_s,i_ to S_s,i+1_. Our specific focus was whether the EEG from the trials S_r,i_ and the subsequent ISI preceding a reversal trials S_r,i+1_ differed from the stability trials S_s,i+1_ and the subsequent ISI.

#### 2.5.1 Global Field Power (GFP)

For each participant, we calculated event-related potentials by separately averaging S_r,i_ and S_r,i+1_ across reversal trials and S_s,i_ and S_s,i+1_ across stability trials. We then used the ERPs to calculate the Global Field Power, “GFP”, which is the spatial standard deviation at a single time point (t) (Lehmann & Skrandies, 1980).

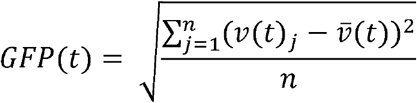

where *v*(*t*)_j_ is the voltage at electrode *j* at time point *t*, 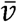 is the mean voltage of all electrodes at time point *t*, and *n* is the total number of electrodes. The GFP is calculated for every time point, resulting in a time series of GFP values.

We were interested in EEG correlates of perceptual destabilization. Given, that it is so far unclear at which time point or at which EEG electrode exactly we should expect such correlates, we used the GFP as a measure that integrates across electrodes and thus reduces the dimensionality of the data. Results from the GFP then allowed us to define a temporal region of interest (ROI) for the next analysis steps.

#### 2.5.2 Data Analysis based on Machine Learning

To investigate how well the EEG data differed between stable and reversal (i.e., destabilized) trials, an Artificial Neural Network (ANN) was trained. For this we used the Deep4Net ANN via the package Braindecode, version 0.5.1 (Schirrmeister et al., 2017), which was specifically designed for the analysis of EEG data. The ANN was trained and evaluated individually for every participant in the following way:

a. From each participant’s trials, 75% were systematically selected for the training and testing of the ANN and the remaining 25% were selected for the evaluation of the ANN. For the training procedure the order of trials was randomized and the reversal trials resampled to match the number of stable trials (the evaluation trials were not randomized or resampled). The data were also all normalized, resulting in all values in a single trial to range between 0 and 1. As a result of the evaluation, for each trial from the evaluation set we either obtained a 1 or 0 as a “*prediction label*” from the ANN classification, indicating either the identification of a stable or destabilized perceptual state. Additionally, for each trial we obtained a binary manual response from the participant indicating either a perceived reversal or perceived stability, which we call “*true label*”. We then calculated *accuracy values* as the number of evaluation trials where the predicted label matched the true label, divided by the total number of trials in the evaluation set.
b. This procedure was executed four times to have each quarter once for the evaluation, resulting in four separate accuracy values in percent discriminability. For each participant, we calculated the median of the resulting four accuracy values. The median accuracy reflects the differentiation between stable and destabilized perceptual states, ideally, indicating how good the destabilization of a perceptual state is reflected in the EEG data. Additionally, and more realistically, the accuracy also reflects how good the ANN was at extracting the relevant information.

One basic idea of this study was, that in the case of endogenous spontaneous perceptual reversals, during observation of an unchanged ambiguous cube stimulus, the EEG contrast described above may reflect a state of perceptual destabilization immediately before the reversal. In contrast, in the case of exogenously induced reversals between the disambiguated cube variants, no such destabilization state should be observed. This should be reflected in a comparison between the ambiguous and disambiguated stimulus conditions.

We applied a bootstrap method to compare distributions of accuracy values for the two conditions in the following way:

(c) We put all the test results from all four test sets together, randomly selected 70 % of these data and calculated an accuracy value from these data. This was done separately for each participant and experimental condition (ambiguous and disambiguated stimuli) 1000 times in a sampling-with-replacement manner.
(d) For each participant we created two accuracy value distributions for the two conditions and compared them with each other by applying a Kolmogorov-Smirnov test.

The analysis steps (a) – (d) were applied for two different temporal ROIs, which were (1) a time window between 300 ms and 700 ms after onset of the stimulus immediately before an indicated perceptual reversal (S_i_) and (2) the subsequent ISI. The choice of the S_i_ time window was based on the results from the previous Global Field Power analysis. Within the time range where the p-values comparing the reversal and stable GFP in S_i_ were below 0.05, a 400 ms time window was chosen to match the time window length of the ISI. The beginning of the time range was determined by finding the first time point where the p-values were less than 0.05 after onset of S_i_ and physiologically plausible (150 ms after stimulus onset (Joos et al., 2020)). This was 304 ms, and therefore, it was rounded down to 300 ms.

##### Estimating Spatial Regions of Interest

The EEG is a rough measure with a relatively low signal-to-noise ratio. Whether a brain signal of interest can be measured on the scalp, depends on a number of factors, like individual brain anatomies, conductivity and thickness of bones and meninges, etc. (e.g., Bach, 1998; Nunez & Srinivasan, 2006). As a consequence, such a signal may be clearly identified in some participants, but be apparently absent in others, even if the neural processes of interest may be highly similar across participants. For this reason, we performed the above-described analysis on the level of individual participants and selected high performers (as defined below) to investigate, which of the 32 EEG electrodes provided the most classification information.

The ANN then was trained and evaluated 8 more times on the high performers and with each training and evaluation a subgroup of electrodes (frontal, central, parietal, occipital, left hemisphere, right hemisphere, parietal left hemispheric, and parietal right hemispheric electrodes) was removed. If the accuracy was lower when removing some electrodes compared to others, this suggested that the missing electrodes contained useful information for the ANN to classify the EEG of S_i_ as a reversal or stable trial.

To quantify the results of these different ANN runs, a ratio between the accuracy values from the Ambiguity Condition and Disambiguation Condition was calculated for each participant. The Disambiguation Condition in this case served as a control as the EEG from the trial immediately before the exogenously induced reversal (S_i_) should have no information about the subsequent reversal (S_i+1_). As a consequence, no indication of perceptual destabilization states should be observed in the data from this condition.

#### 2.5.3 Source Analysis

Standard EEG electrode positions were assumed for all q=32 electrodes according to the 10-20 system since no individual EEG electrode positions from the individual participants were available from the dataset. We used the “fsavagerage” (Fischl et al., 1999) template T1 image as provided by the Freesurfer image analysis suite (https://surfer.nmr.mgh.harvard.edu/). The forward model was computed using the boundary element method (BEM, Fuchs et al., 2002) as provided by MNE-Python (Gramfort, 2013). Each shell (brain, skull, and scalp tissue) was composed of 5120 vertices. Conductivity was set to 0.3 S/m for brain and scalp tissue, and 0.06 S/m for the skull.

The source space was created using p=1284 dipoles with icosahedral spacing, i.e. dipoles were placed along the cortical folding by iterative subdivision of a icosahedron (cf. https://mne.tools/, Gramfort et al., 2013). In order to reduce computational complexity and based on the reasonable physiological assumption, a fixed orientation of dipoles orthogonal to the surface of the cortical sheet was assumed.

Inverse solutions were calculated for the ERPs spanning 300 to 700 ms after the onset of S_i_. This was the same range used for the classification of the ANN as described above. In order to mitigate the problem of missing individual forward models, we adapted a group-inversion scheme similar to the one described by Friston et al. (2015).

As a first step, we concatenated the reversal- and stability-ERPs of all participants. Next, we identified a global set of active sources by calculating the flexible multi-signal classification (FLEX-MUSIC) inverse solution on the concatenated ERPs (Hecker et al., 2023). FLEX-MUSIC is a recently developed improvement of the well-established recursive MUSIC approach to solve EEG inverse problems, that can accurately estimate not only the location but also the spatial extent of neural sources. Finally, weighted minimum norm estimates (wMNE, (Pascual-Marqui, 1999)) were calculated on the global set of active sources for each participant and condition.

## 3 Results

### 3.1 Global Field Power shows differences in conditions already in the trial before the reversal

The GFP of reversal- and stable-ERPs, together with the GFP difference traces (reversal GFP minus stability GFP) are shown in Figure 3. From this Figure, it is clear that in the Disambiguation Condition (red traces, left), the activity in S_r,i_ is similar to that of S_s,i_. As a consequence, the difference GFP trace in the S_i_ time window is close to zero. The GFP difference trace in the S_i+1_ time window, however, shows a highly significant deviation from the zero line, indicating a clear GFP difference between a reversed and a stable percept (maximal significance at 427 ms after onset if S_i+1_, p=1.6*10^−8^, Cohen’s d=2.19).

**Figure 3.**
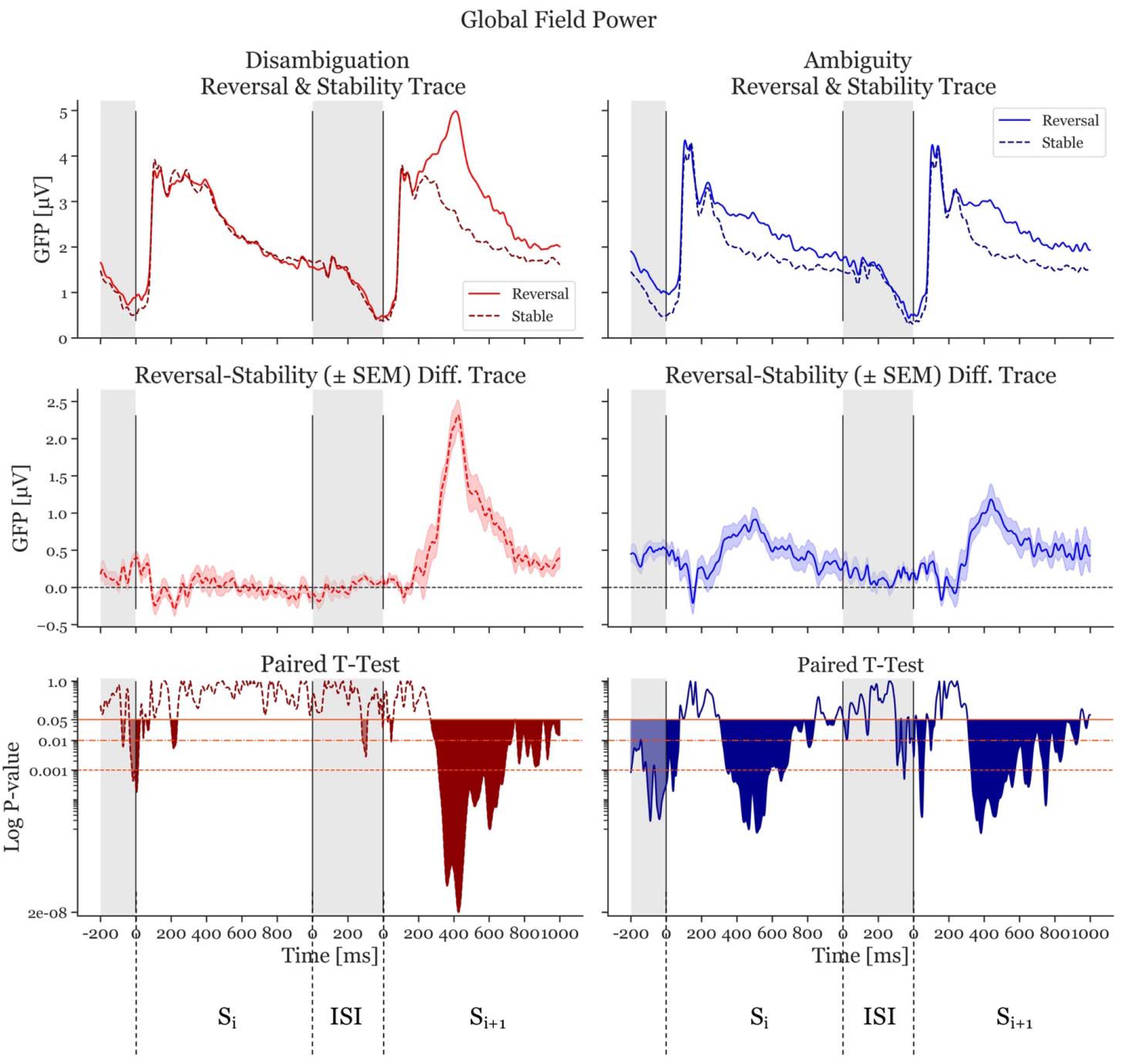
Global Field Power (GFP) and p-values of the Disambiguation (red) and Ambiguous (blue) Conditions. The two top panels show the GFP averaged across participants. The dashed, darker traces depict the stable condition, whereas the lighter, continuous trace depicts the reversal condition. The middle two panels depict difference GFP traces of the top row (reversal minus stability) resulting in the difference traces. The shaded area is ± standard error of the mean. The bottom panels shows the p-values logarithmically scaled. The orange-red horizontal lines depict alpha values of 0.05, 0.01, and 0.001, respectively. The filled (red and blue) areas indicate statistically significant time periods. The gray areas indicate interstimulus interval time ranges. The first range shown is the ISI preceding S_i_. The next 1000 ms represent the onset period of S_i_ and the 1000 ms after the interstimulus interval show S_i+1_.

The Ambiguity Condition (Fig. 3, blue traces, right) presents a different pattern. The difference GFP trace in S_i_ shows a significant deviation from zero, with a similar shape to the trace from S_i+1._ The most significant effect in the S_i_ time window is at 514 ms after stimulus onset (p = 7.59*10^−6^, Cohen’s d = 1.35). The most significant effect in the S_i+1_ time window is at 382 ms after stimulus onset (p =7.3*10^−6^, Cohen’s d=1.06).

### 3.2 ANN uses parietal information to classify reversal and stable trials already approximately 1000 ms before an upcoming reversal

First, the ANN was trained and evaluated on the most statistically significant 400 ms of the stimulus presentation window S_i_ based on the GFP results (i.e., 300-700 ms after stimulus onset, cf. Figure 3). This resulted in a mean accuracy of 59.22% for the Ambiguity Condition and 52.29% for the Disambiguation Condition. Figure 4 indicates that all but two participants had a higher accuracy in the Ambiguous Condition compared to the Disambiguation Condition.

**Figure 4.**
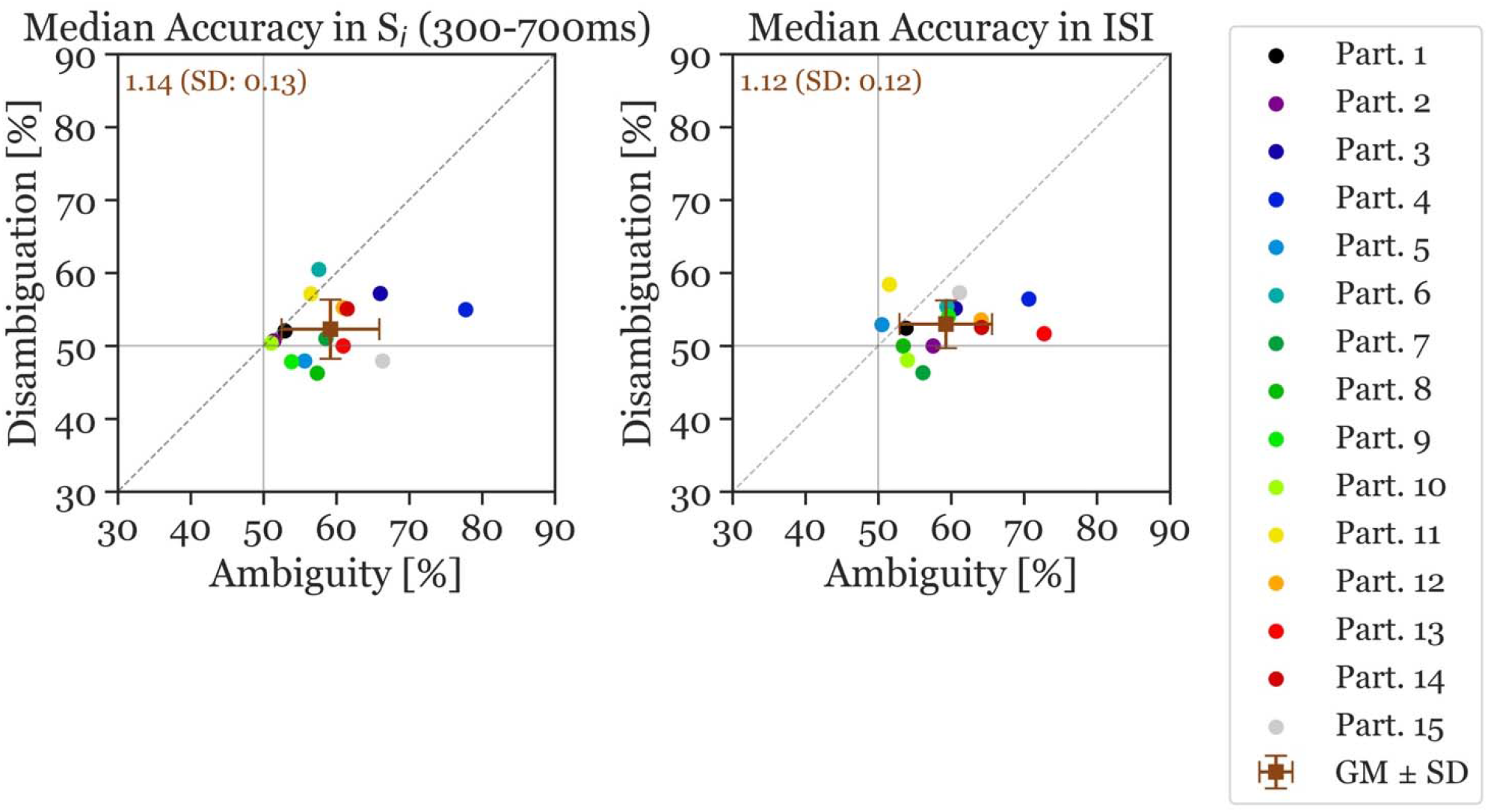
Median accuracy of individual participants (colored circles) and the grand mean ± standard deviation (brown squares). The x-axis shows the accuracy in the Ambiguous Condition and the y-axis the accuracy in the Disambiguation Condition. The left panel shows the accuracy calculated during the S_i_ Stimulus Time Window and the right panel shows the accuracy during the ISI Time Window (reversal and stable). The number inserted in the top left shows the mean accuracy ratio (ambiguous divided by disambiguated) accuracies (with the standard deviation).

Next, the ANN was trained and evaluated on the 400 ms ISI, resulting in a mean accuracy across all participants of 59.26% for Ambiguous and 53% for Disambiguated Conditions. Again, all but the same two participants as mentioned previously had a higher accuracy in the Ambiguous Condition during the ISI. The average accuracy ratio between ambiguous and disambiguated is slightly larger in S_i_ (1.14) compared to during the ISI (1.12). The standard deviation of the accuracy ratios is also larger in S_i_ (0.13) compared to during the ISI (0.12). Moreover, the accuracies show more variance in the Ambiguity Condition than in the Disambiguation Condition.

The ambiguous and disambiguation accuracy distributions were compared, in both the 300-700 ms in S_i_ and the 400 ms of the ISI, using the Kolmogorov-Smirnov test. The Kolmogorov-Smirnov test was used to quantify the difference between the accuracy distributions of the Ambiguous and Disambiguated condition for each participant. Fig. 5 displays distributions based on bootstrapping from three representative example participants together with the related test results. The three examples indicate the strong differences in the discriminatory power of the EEG between participants. The Kolmogorov-Smirnov test results from all participants is shown in Figure 6. One interesting aspect in this graph is the linear relation between the ISI and S_i_ data, indicating that if a participant’s EEG showed a strong discriminatory power in the S_i_ time window, it was also relatively strong in the subsequent ISI. The correlation between the statistic values of these two time windows was ρ=0.8 (p=0.0009). Furthermore, 60 % of participants showed larger discriminatory power in the S_i_ time window compared to the ISI time window. Finally, the graph identifies participants 2, 3, 4, 9, 12, 13, 14, and 15 as high performers. Participant 13 has more discriminatory EEG power in the ISI compared to S_i_, while for the others there is more discriminatory power in S_i_ compared to the ISI. The high performers were determined by choosing all participants that had KS a statistic score larger than or equal to 0.8 in either the S_i_ time window or the ISI time window.

**Figure 5.**
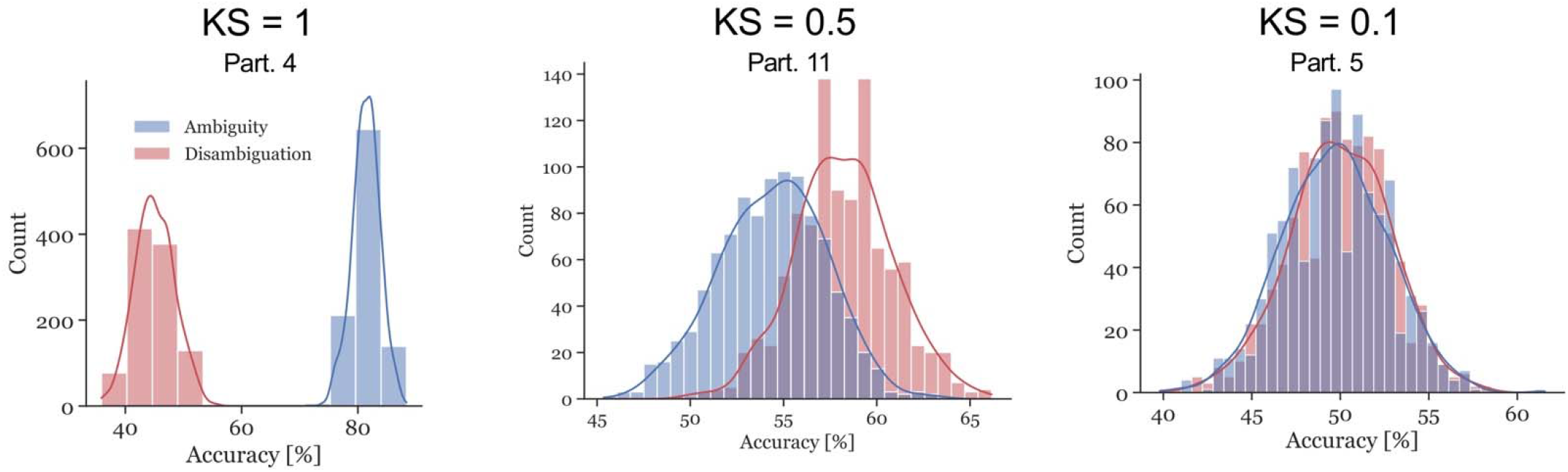
Examples of accuracy distributions resulting from the bootstrap method and the corresponding Kolmogorov–Smirnov (KS) test statistic for participants 4, 11, and 5. The red distributions represent the Disambiguation Condition accuracies, the blue distributions represent the accuracies in the Ambiguous Condition. The higher the KS test statistic, the further apart the distributions are.

**Figure 6.**
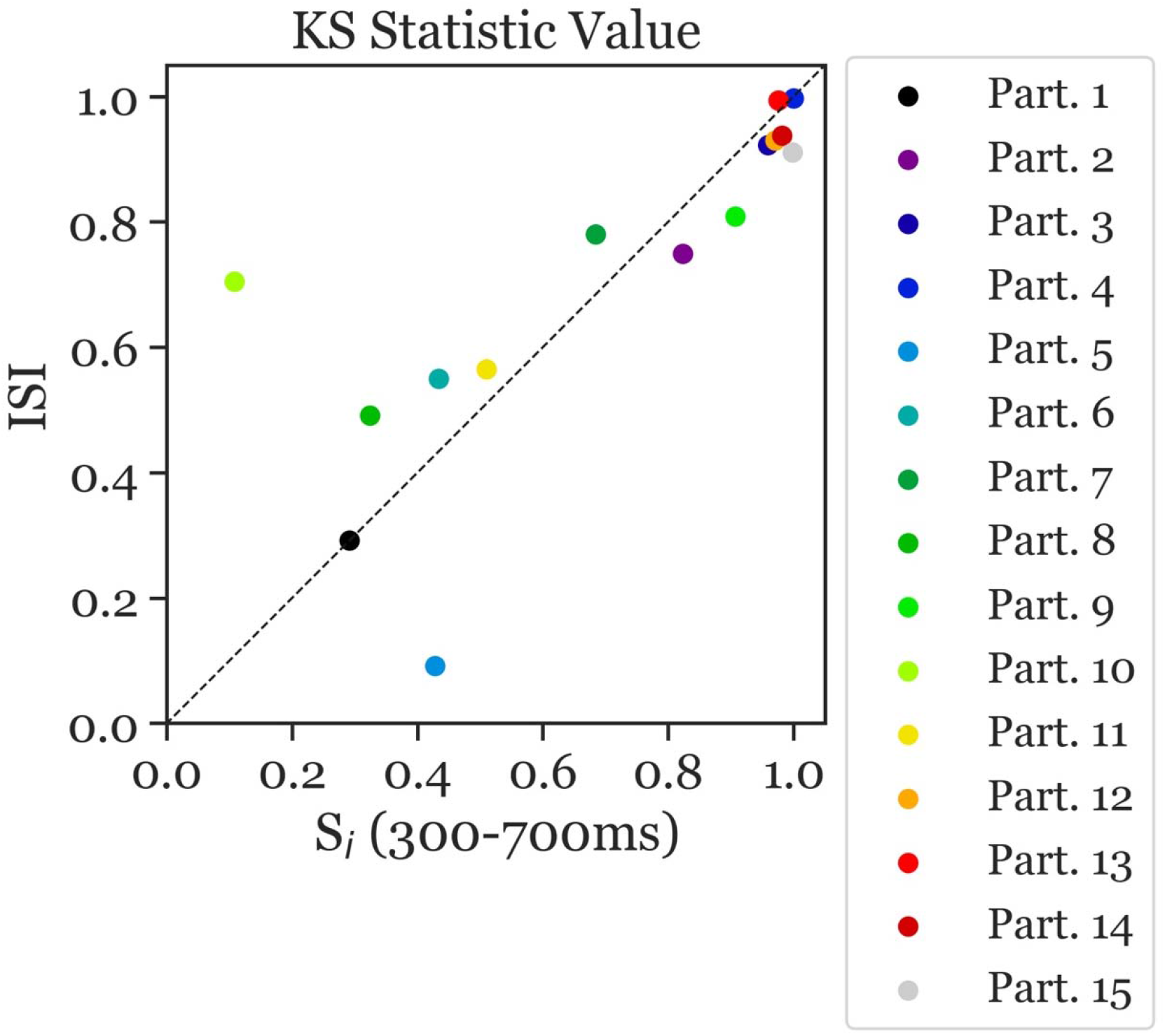
Kolmogorov–Smirnov (KS) test statistic values indicating the separability of the distribution of ambiguous and disambiguated accuracies in S_i_ (300-700ms) and the ISI. The x-axis presents the KS statistic of S_i_ Time Window. The y-axis presents the statistic of the ISI Time Window. The colors correspond to the individual participants. Most participants with high/low separability in the S_i_ Time Window also showed high/low values in the ISI Time Window, with overall better separability in the S_i_ Time Window than in the ISI Time Window (i.e., more data points below the diagonal). Moreover, largest separability can be observed in participants 2, 3, 4, 9, 12, 13, 14, and 15.

In the following spatial analysis steps, we focused on these 8 high performers. We repeated the ANN calculations with different subsets of electrodes in order to identify electrode subsets that were necessary to obtain high discrimination performance.

Figure 7 shows the results of this analysis. Without the removal of any electrodes, our high-performance participants had an average accuracy ratio of 1.19 (SD=0.13). When the parietal electrodes were removed, the median accuracy was slightly decreased (accuracy ratio=1.18; SD=0.12). Remarkably, for all other variants of electrode removal, the accuracy increased. Moreover, removing the left parietal electrodes increases the accuracy slightly more than removing the right parietal electrodes (accuracy ratio=1.25; SD=0.095 versus accuracy ratio=1.22; SD=0.098). Additionally, removing all electrodes from the left hemisphere increased the accuracy slightly more than removing the right hemisphere electrodes (accuracy ratio=1.30; SD=0.12 vs accuracy ratio=1.29; SD=0.16). Overall, removing the left-hemisphere electrodes provides the highest accuracy ratio among all variants.

**Figure 7.**
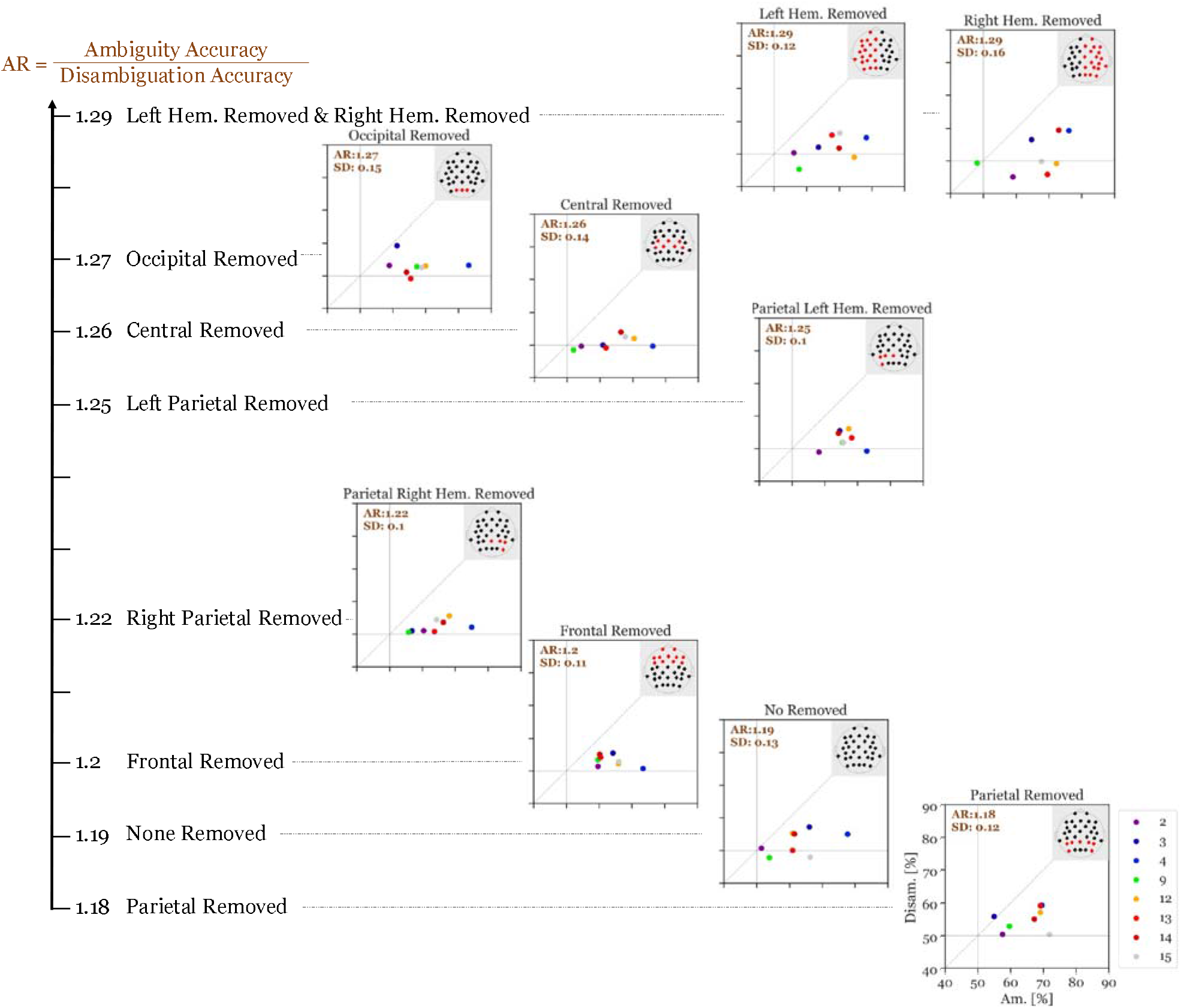
Accuracy ratio of S_i_ (300-700ms after stimulus onset) of the top 8 performers with different electrode subsets removed with each run. The values show the mean accuracy ratio of all top performers. The different colors depict the different participants and the numbers in the top left of each panel represents the mean accuracy ratio (Ambiguity divided by Disambiguation; with the standard deviation). The larger the distances (to the right and to the top) of the individual icons from the horizontal and vertical grey lines are, the higher is the discriminatory power of the respective participant’s EEG data.

### 3.3 Source Localization

Source localization was performed in a two-stage procedure. First, the global set of active sources was identified with the EEG source analysis method FLEX-MUSIC, then the neural activity at these regions was estimated on the level of individuals participants.

Notably, a relatively confined set of active regions was found (cf. Figure 8), which is remarkable given the presumably high inter-individual variance of the EEG data. The results of the source localization suggest that the parahippocampal place area (PPA) most significantly differs between reversal and stable trials (p=3.96*10^−05^; cf. Figure 8 and Table 1). A summary of the brain areas and the corresponding t- and p-values is shown in Table 1. Other regions that significantly differ are the fusiform gyrus, the lingual gyrus, the entorhinal cortex, the posterior cingulate cortex, and the precuneus. Most activity seems to be located in the right hemisphere.

**Figure 8.**
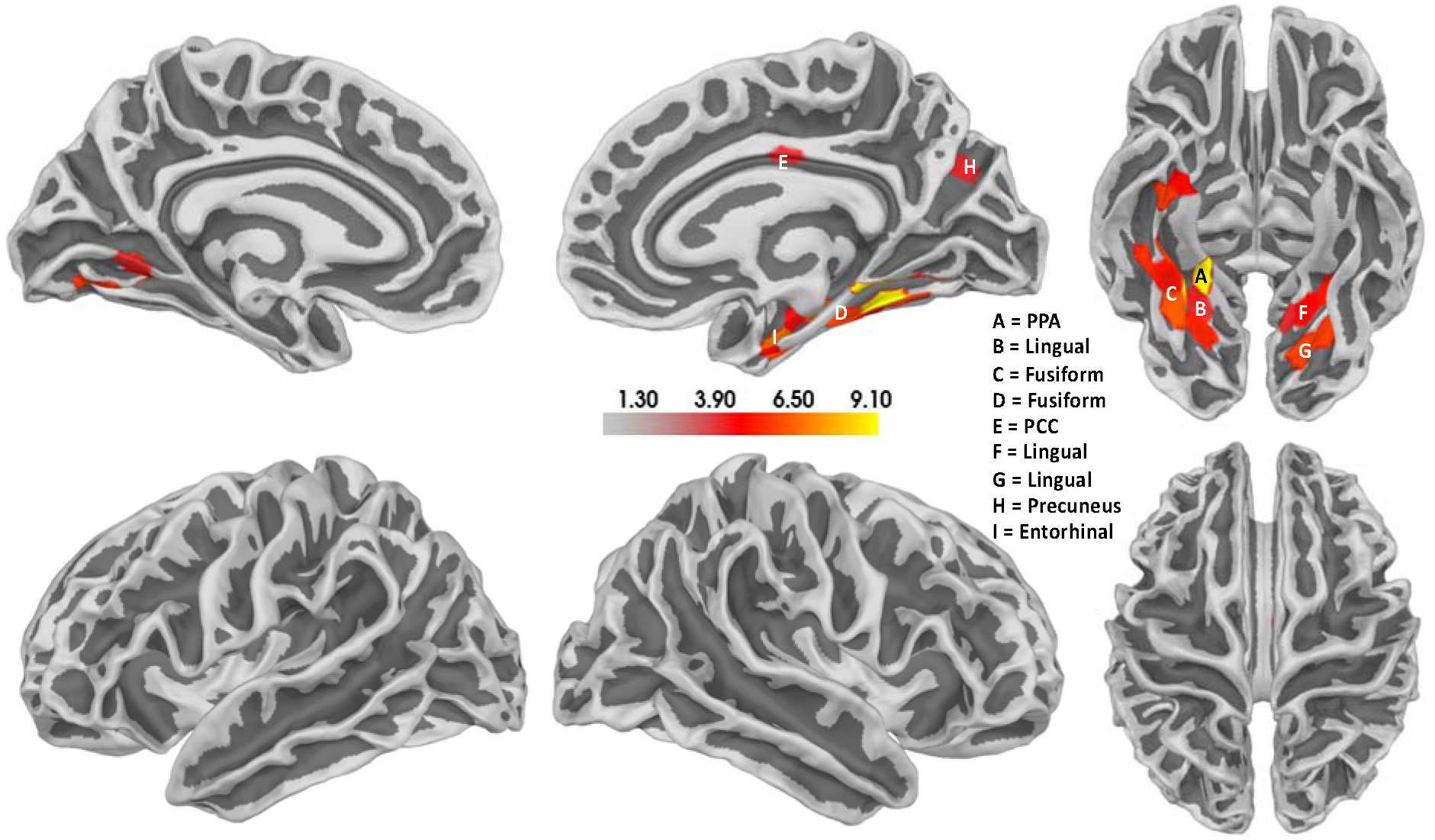
Results of the source localization with thresholded t-values for the Ambiguity Condition. Reversal ERPs compared to stable ERPs. The size of the cluster does not represent the relevance of the cluster. Activity seems to be concentrated mainly in the right hemisphere, specifically the parahippocampal place area (PPA). There also seems to be some activity in the left hemisphere in a similar area. Additionally, the posterior cingulate cortex (PCC) also seems to be active.

**Table 1.** The location of all significant voxels that are in named brain areas in order of most significant to least significant.

| Brain Area | Hemisphere | Legend in Figure 8 | T | p |
| --- | --- | --- | --- | --- |
| Parahippocampal place area | Right | A | 9.1 | $3.96*10^{-05}$ |
| Fusiform gyrus | Right | C | 9.1 | $4.09*10^{-05}$ |
| Lingual gyrus | Left | G | 6.34 | 0.0004 |
| Fusiform gyrus | Right | C | 6.12 | 0.0005 |
| Lingual gyrus | Left | F | 5.74 | 0.0007 |
| Fusiform gyrus | Right | D | 5.71 | 0.0007 |
| Fusiform gyrus | Right | D | 5.68 | 0.0008 |
| Lingual gyrus | Left | G | 5.64 | 0.0008 |
| Fusiform gyrus | Right | C | 5.37 | 0.001 |
| Lingual gyrus | Right | B | 5.22 | 0.001 |
| Lingual gyrus | Left | F | 5.98 | 0.002 |
| Fusiform gyrus | Right | B | 4.93 | 0.002 |
| Entorhinal cortex | Right | I | 4.67 | 0.002 |
| Lingual gyrus | Left | F | 4.65 | 0.002 |
| Posterior cingulate cortex | Right | E | 3.82 | 0.007 |
| Lingual gyrus | Right | B | 3.81 | 0.007 |
| Lingual gyrus | Left | F | 3.79 | 0.007 |
| Precuneus | Right | H | 3.73 | 0.007 |

## 4 Discussion

The primary aim of the present study was to identify EEG correlates of a destabilized perceptual brain state preceding a spontaneous perceptual reversal of an ambiguous Necker lattice. Given the absence of spatial and temporal regions of interest from the EEG literature to focus this question on, we started with a Global Field Power (GFP) analysis. We compared the GFP from time periods during perceptual stability with those about 1 second before a perceptual reversal and found large GFP effects with ambiguous lattice stimuli. As expected, no such effects were observed with the disambiguated control lattice stimuli.

The concept of perceptual destabilization is not well established in the literature. It is, thus, unclear, how common the underlying neural processes are in time and brain areas across participants. We, therefore, focused the analysis steps subsequent to the GFP analysis on the level of individual participants using ANNs. We determined the accuracy of the individually trained ANNs to discriminate between stable and destabilized brain states. This analysis identified eight (out of 15) participants with high EEG discrimination performance already about one second before an upcoming reversal. By repeated re-analysis of the data from these eight participants with the systematic removal of subsets of electrodes, we were able to identify the parietal cortex as a critical brain area for the prediction of an upcoming reversal. A subsequent source localization analysis revealed the parahippocampal place area (PPA) as the most statistically significant brain area for reversal prediction. Previous studies identified a network of frontal and parietal regions being active and relevant during spontaneous perceptual reversals (J. Brascamp et al., 2018; Watanabe, 2021).

Interestingly, neither our ANN analysis nor the source analysis indicated the relevance of this network for destabilization. Needless to say that the source localization needs to be taken with caution, due to the missing individual forward models.

### 4.1 Potential Limitations

#### 4.1.1 Limited number of trials and participants

In the present study, we measured 21 participants. Six had to be excluded due to too low reversal rates (see Methods), resulting in a dataset of 15 participants. This is not a very large sample to make strong claims on. On the other hand, the majority of our analyses was realized on the level of individual participants. From our 15 analysed participants, we identified eight high performers. We cannot say whether the effects we identified in 53% of our participants can also be identified in 53% of the population. However, we regard our results as a good starting point for follow-up studies with a specific focus on perceptual representations and their stability criteria. Further below, we will also suggest possible next steps to increase the discrimination accuracies.

#### 4.1.2 Are perceptual reversals during discontinuous stimulus presentation a good model for the continuous case?

The Introduction Section contains an entire paragraph on the question about how to measure spontaneous endogenous perceptual reversals of an ambiguous stimulus with high temporal resolution. In the present study, we used the Onset-Paradigm as a possible way to address this principal problem. One can now ask whether a changed percept from one stimulus presentation to the next is really comparable with a spontaneous reversal during a continuous observation of the ambiguous stimulus?

We think “yes”, as we have already discussed this question in detail in our review paper in 2012 (Kornmeier & Bach, 2012). In short, continuous observation is never really continuous, because of around 15 eye blinks per minute on average (Sforza et al., 2008), with a duration of around 200 ms (Caffier et al., 2003) and the ensuing saccadic suppression (Binda & Morrone, 2018). Moreover, a number of behavioral studies showed a continuous transition of reversal rates from continuous stimulus presentation to short reoccurring interruptions up to 400 ms duration (Kornmeier et al., 2007; Orbach et al., 1963, 1966). Clearly, however, interrupted stimulus presentation is not the same as continuous presentation. Based on the present results, it may thus be an interesting next step to train ANNs with EEG data from discontinuous presentation and apply the trained ANN to EEG data from continuous observation.

### 4.2 Why do we only see clear discrimination accuracy in 53% of the participants?

The scalp EEG is a relatively rough measure. Whether a neural signal reaches the scalp and can be detected by the scalp EEG electrodes depends, amongst other factors, on the individual folding of the brain. The activity of neighboring brain sources of activity can be superimposed and weaken each other. Furthermore, neural signals related to a certain processing step are typically superimposed on background activity, not necessarily related to this processing, but affecting the signal-to-noise ratio of the signal in question (Arieli et al., 1996; Bach, 1998). These and other factors contribute to a large inter-individual variability of brain activity, even if participants observe the same stimulus or execute the same task (e.g. Kornmeier et al., 2014). An impressive demonstration of the inter-individual variability during observation of multistable stimuli has recently been presented by Wexler (2018). A huge number of reported EEG effects, including work from our lab, resulted from group statistics and were difficult to observe in a large number of individual participants (e.g. Ehm et al., 2011). Hence, it is not surprising that in the present data some participants showed weak or even no effects. The low signal-to-noise ratio of single-trial EEG may also explain that even for the eight high performers the discrimination accuracy remained below 80%. Finally, another potential limitation could be the low number of trials the ANN had available for training. Due to potential fatigue effects in our participants, typically resulting in rising numbers of body movement artifacts, wrong key presses etc., it was difficult to increase the number of trials to be enough for training the ANN.

Beyond these rather technical reasons, another interesting more functional factor may play a role. Polgári et al. (2020) recently published an interesting study about eye-movements during observation of the Necker cube. In the present study, participants were instructed to fixate on a fixation cross in the center of the cube stimuli in order to minimize eye-movement artifacts in the EEG. In Polgári et al.’s study participants were allowed to freely move their eyes. The authors analysed fixations and clustered them based on the horizontal fixation location as an either right or left fixation cluster with respect to the vertical midline of the visual field in which the Necker cube was presented centrally.

They found that in most cases, in which participants reported a reversal, the eyes also moved from one cluster to the other. However, they recorded about twice as many changes of the horizontal eye position between clusters than reversals, which means that not every cluster change was accompanied by a clear and conscious perceptual reversal. Polgári et al.’s explanation of this observation is in contrast to the basic assumption underlying the present study. We *a priori* assumed that participants’ percepts become stable immediately after a reversal and only slowly destabilized towards the next perceptual reversal. Comparing the EEG from the middle of this proposed temporal window of perceptual stability with the EEG close to a reversal should therefore reveal potential EEG correlates of perceptual stability state differences (stable versus destabilized). Polgári et al., in contrast, postulated that the perceptual brain state can also destabilize in between two button presses, but sometimes restabilizes back to the perceptual interpretation where it started. They further speculate that such unconscious destabilizations may come with a partial or total departure from one of the clustered horizontal eye positions, which would explain the larger number of cluster changes than button presses. Interestingly, indications of unconscious destabilizations during observation of ambiguous figures have also been reported by other groups (e.g., Pastukhov & Braun, 2007; Pastukhov & Klanke, 2016).

As described above, our discrimination accuracy measure is based on a postulated EEG contrast between perceptually stable and unstable brain states. If Polgári et al.’s postulate of unconscious destabilizations is correct, it may be possible that the time windows we selected and labelled as perceptually stable brain states between two button presses may contain periods of unconscious destabilizations. This may lead to a larger variability and a smaller signal-to-noise ratio of our measure. Accepting Polgári et al.’s interpretation, horizontal eye-movement data can be used in a future replication of the present study to better determine perceptually stable brain states.

### 4.3 What can the current results contribute to the understanding of perceptual representations?

When our participants look at a black screen and suddenly a white lattice stimulus occurs, a clear and distinct pattern of EEG activity can be identified within the first 600 – 800 ms after stimulus onset.

This pattern typically has a temporal profile which can be observed, e.g., in the ERP with stimulus onset as the reference time point. In the first 300 ms we see so-called exogenous signatures with typically sharp and relatively large deviations from baseline, followed by broader components, like the P300-family (e.g., Fig. 2c in Kornmeier & Bach, 2006). These signatures in the ERP (lower-frequency range) are typically accompanied by modulations in higher frequencies (Ehm et al., 2011). These signatures are typically discussed as processing steps along a hierarchy of perceptual processing on the way to create a stable perceptual interpretation (Dehaene & Changeux, 2011; Kornmeier & Bach, 2012; Sergent et al., 2017; VanRullen & Thorpe, 2002). Interestingly, after the percept has been established and is stable, the measurable signals seem to fade out, raising the fundamental question of how stable perceptual interpretations are represented in the brain over time (e.g., Krüger, 2022). Of course, given the quality of the EEG, we cannot infer that the absence of a measurable signal means no processing in the brain. At this point, ANNs become relevant, because these methods seem to be more sensitive to subtle signals.

Given that no precise spatio-temporal regions of interest were available for our experimental question and taking into account, that the signatures we were looking for, could be highly variable in space and time across observers and at the same time very small, we decided to focus on within-participant statistics and apply ANN methods for our analysis after a group GFP analysis. In the GFP analysis of the Ambiguity Condition we found surprisingly large effects in a time window already about one second before a reversed percept was established. Two observations are of particular interest here:

1. In our analysis, we focused on two time windows of interest, the S_i_ time window, where the last lattice stimulus before the perceptual reversal was presented, and the subsequent ISI. In the GFP analysis we found a relatively long time period (about 400 ms) of significance with huge effect sizes up to 1.35. We also found significant effects in the subsequent ISI, but with much shorter time periods and smaller effect sizes. Interestingly, while the GFP analysis indicates that most of the destabilization information seems to necessitate the presence of a stimulus (S_i_ time window), the subsequent ANN analysis draws a more sophisticated picture. In contrast to the GFP results, Figure 6 indicates that our top performers showed comparable discriminatory power in both the S_i_ time window and in the subsequent ISI. This supports the observation that the ANN methods are more sensitive. Moreover, it indicates that the destabilization state of the perceptual system can also be read out from EEG data in the absence of a visual stimulus.
2. During the S_i_ presentation time window, the period of high significance started at about 300 ms after stimulus onset. Interestingly, 300 ms are discussed as an estimate of time, necessary for a perceptual interpretation to become conscious, after the early visual processing steps have finished (Atmanspacher et al., 2008; Atmanspacher & Filk, 2013; Dehaene & Changeux, 2011; Kornmeier, Friedel, et al., 2017; Kornmeier & Bach, 2012). If we interpret our observed effects as correlates of perceptual destabilization, then the GFP analysis results indicate, that the early visual processing units are less relevant. This hypothesis is further supported by the observation that eliminating the occipital electrodes did not substantially reduce the discriminatory power (cf. Fig. 7)

Our current results do not allow strong conclusions about the ‘nature of a perceptual representation’, nor on the definition of precise spatial regions of interest for subsequent studies. However, the huge effect sizes in precisely defined time windows make this study a perfect starting point for subsequent studies about stable and unstable perceptual brain states.

### 4.4 Conclusions

We (Ehm et al., 2011) and others (Britz et al., 2009, 2010; Nakatani & van Leeuwen, 2013) already reported about EEG effects shortly before a reversed percept of an ambiguous Necker lattice is established. Our present results indicate that such anticipatory activity is already present at least one second before the reversal. Technical reasons (not enough single trials and number of reversals), unfortunately, did not allow us to go further back in time. It is, thus, possible that EEG indicators of destabilization and, thereby, of an upcoming perceptual reversal are present earlier. It would be highly interesting to see whether this is really the case.

In his Principles of Psychology, the great psychologist William James wrote about stable mental representations and transient, unstable states between them (James, 1890). He emphasized that the stable states are the “substantive parts” that enter consciousness, while the “transitive parts” are typically very fast and stay unconscious. The basic idea of this has further been elaborated in terms of categorical versus acategorical perceptual/mental states (Atmanspacher, 1992; Atmanspacher & Fach, 2019; Feil & Atmanspacher, 2010).

The present findings indicate that the perceptual system is already in an unstable state about one second before a perceptual reversal. The eye-tracking data from Polgári et al. (2020) further indicate that transient periods of perceptual instability may occur between two consciously perceived reversal events. All of this indicates that what James described as the transient state may only be the tip of the iceberg, i.e. the point of maximal instability (Kornmeier & Bach, 2012) and that perceptual instability is a gradual and longer lasting phenomenon rather than being binary and short.

We demonstrated that perceptual instability can be measured with the EEG. Moreover, the effects are large enough to be visible in a considerable number of individual participants. Taking into account the findings from Polgàri et al., it may even be possible to increase the sensitivity of our measures by eliminating time windows of apparent stability, in which eye tracking data, however, would indicate destabilization. This might make some of the low performing participants become high performers.

We interpret the present EEG effects as a measure of perceptual instability, i.e. as reflecting the difference in neural activity between stable and destabilized perceptual states. An interesting question for subsequent studies now, is to see whether this measure reflects perceptual (in)stability on a continuous scale, i.e. whether the size of the effect reflects the amount of instability of the system.

Furthermore, a number of studies show deviating perceptual reversal dynamics, when participants with psychiatric disorders observe ambiguous stimuli (e.g. Kornmeier, Wörner, et al., 2017; Notredame et al., 2014; Robertson et al., 2016; Schmack et al., 2015). These deviating perceptual dynamics in patients may indicate an underlying imbalance between stable and unstable brain states. Deviating patterns of EEG variability and other stability measures of brain activity further confirm these observations (Foerster et al., 2021; Hecker et al., 2022; Huang et al., 2022; Koshiyama et al., 2018; Marques-Carneiro et al., 2021). In summary, the present paradigm together with the identified physiological effects are a perfect starting point for further research about stability features of brain activity comparing patients with healthy controls. As a result, our measure of perceptual instability may become a realistic candidate in the urgent search for biomarkers of psychiatric disorders.

## 5 Conflict of Interest

The authors declare that the research was conducted in the absence of any commercial or financial relationships that could be construed as a potential conflict of interest.

## 6 Author Contributions

EJ, LH, and JK contributed to the conception and design of the study. EJ and LH contributed to the data acquisition. MW, LH, JK and AA contributed to the conception of the analysis. MW and LH contributed to the analysis of the data. All code was written by MW and LH. JK and MW wrote the first draft of the manuscript. LTvE and JK contributed to the acquisition of the funding. All authors contributed to manuscript revision, read, and approved the submitted version.

## 7 Funding

The project was funded by Neurex, the Deutsch-Franzoesische Hochschule (DFH), European Campus (Eucor) seeding money, and the Institute for Frontier Areas of Psychology and Mental Health (IGPP).

## 8 Acknowledgments

We thank Julia Schipp for the data acquisition.

## Notes

### Competing Interest Statement

The authors have declared no competing interest.

### Summary of Updates

Formatting of figures in PDF.

## References

Abdallah, D., & Brooks, J. L. (2020). Response dependence of reversal□related ERP components in perception of ambiguous figures. Psychophysiology, 57(12). https://doi.org/10.1111/psyp.13685

American Clinical Neurophysiology Society. (2006). Guideline 5: Guidelines for standard electrode position nomenclature. J Clin Neurophysiol, 23(2), 107–110.

Arieli, A., Sterkin, A., Grinvald, A., & Aertsen, A. (1996). Dynamics of Ongoing Activity: Explanation of the Large Variability in Evoked Cortical Responses. Science, 273(5283), 1868–1871. https://doi.org/10.1126/science.273.5283.1868

Atmanspacher, H. (1992). Categoreal and acategoreal representation of knowledge. Cogn Sys, 3, 259–288.

Atmanspacher, H., Bach, M., Filk, T., Kornmeier, J., & Römer, H. (2008). Cognitive Time Scales in a Necker-Zeno Model for Bistable Perception. The Open Cybernetics and Systemics Journal, 2, 234– 251.

Atmanspacher, H., & Fach, W. (2019). Exceptional Experiences of Stable and Unstable Mental States, Understood from a Dual-Aspect Point of View. Philosophies, 4(1), 7. https://doi.org/10.3390/philosophies4010007

Atmanspacher, H., & Filk, T. (2013). The Necker-Zeno Model for Bistable Perception. Topics in Cognitive Science, 800–817. https://doi.org/10.1111/tops.12044

Bach, M. (1998). Electroencephalogram (EEG). In G. K. von Schulthess & J. Hennig (Eds.), Functional Imaging (pp. 391–408). Lippincott-Raven.

Bach, M. (2018). Homepage of the Freiburg Visual Acuity & Contrast Test (‘FrACT’). http://www.michaelbach.de/fract.html

Bartels, A. (2021). Consciousness: What is the role of prefrontal cortex? Current Biology, 31(13), R853–R856. https://doi.org/10.1016/j.cub.2021.05.012

Basar-Eroglu, C., Struber, D., Kruse, P., Basar, E., & Stadler, M. (1996). Frontal gamma-band enhancement during multistable visual perception. Int J Psychophysiol, 24(1–2), 113–125.

Binda, P., & Morrone, M. C. (2018). Vision During Saccadic Eye Movements. Annual Review of Vision Science, 4(1), 193–213. https://doi.org/10.1146/annurev-vision-091517-034317

Blake, R. (2001). A Primer on Binocular Rivalry, Including Current Controversies. Brain and Mind, 2, 5–38.

Blake, R., Brascamp, J., & Heeger, D. J. (2014). Can binocular rivalry reveal neural correlates of consciousness? Philosophical Transactions of the Royal Society B: Biological Sciences, 369(1641), 20130211–20130211. https://doi.org/10.1098/rstb.2013.0211

Blake, R., & Logothetis, N. K. (2002). Visual competition. Nature Rev Neurosci, 3(1), 13–21. https://doi.org/10.1038/nrn701

Brascamp, J., Sterzer, P., Blake, R., & Knapen, T. (2018). Multistable Perception and the Role of the Frontoparietal Cortex in Perceptual Inference. Annual Review of Psychology, 69, 77–103. https://doi.org/10.1146/annurev-psych-010417-085944

Brascamp, J. W., de Hollander, G., Wertheimer, M. D., DePew, A. N., & Knapen, T. (2021). Separable pupillary signatures of perception and action during perceptual multistability. ELife, 10, e66161. https://doi.org/10.7554/eLife.66161

Britz, J., Landis, T., & Michel, C. M. (2009). Right parietal brain activity precedes perceptual alternation of bistable stimuli. Cereb Cortex, 19(1), 55–65.

Britz, J., Pitts, M. A., & Michel, C. M. (2010). Right parietal brain activity precedes perceptual alternation during binocular rivalry. Hum Brain Mapp, 32(9), 1432–1442. https://doi.org/10.1002/hbm.21117

Caffier, P. P., Erdmann, U., & Ullsperger, P. (2003). Experimental evaluation of eye-blink parameters as a drowsiness measure. Eur J Appl Physiol, 89(3–4), 319–325. https://doi.org/10.1007/s00421-003-0807-5

Carter, O., Konkle, T., Wang, Q., Hayward, V., & Moore, C. (2008). Tactile Rivalry Demonstrated with an Ambiguous Apparent-Motion Quartet. Current Biology, 18(14), 1050–1054. https://doi.org/10.1016/j.cub.2008.06.027

Conrad, V., Vitello, M. P., & Noppeney, U. (2012). Interactions between apparent motion rivalry in vision and touch. Psychol Sci, 23(8), 940–948. https://doi.org/10.1177/0956797612438735

Crick, F., & Koch, C. (1998). Consciousness and neuroscience. Cereb Cortex, 8(2), 97–107.

Darki, F., & Rankin, J. (2020). Perceptual rivalry with vibrotactile stimuli [Preprint]. Neuroscience. https://doi.org/10.1101/2020.09.17.301358

de Jong, M. C., Vansteensel, M. J., van Ee, R., Leijten, F. S. S., Ramsey, N. F., Dijkerman, H. C., Dumoulin, S. O., & Knapen, T. (2020). Intracranial Recordings Reveal Unique Shape and Timing of Responses in Human Visual Cortex during Illusory Visual Events. Current Biology, 30(16), 3089–3100.e4. https://doi.org/10.1016/j.cub.2020.05.082

Dehaene, S., & Changeux, J. P. (2011). Experimental and theoretical approaches to conscious processing. Neuron, 70(2), 200–227. https://doi.org/10.1016/j.neuron.2011.03.018

Devia, C., Concha-Miranda, M., & Rodríguez, E. (2022). Bi-Stable Perception: Self-Coordinating Brain Regions to Make-Up the Mind. Frontiers in Neuroscience, 15, 805690. https://doi.org/10.3389/fnins.2021.805690

Duman, I., Ehmann, I. S., Gonsalves, A. R., Gültekin, Z., Van den Berckt, J., & van Leeuwen, C. (2022). The No-Report Paradigm: A Revolution in Consciousness Research? Frontiers in Human Neuroscience, 16, 861517. https://doi.org/10.3389/fnhum.2022.861517

Ehm, W., Bach, M., & Kornmeier, J. (2011). Ambiguous figures and binding: EEG frequency modulations during multistable perception. Psychophysiology, 48(4), 547–558. https://doi.org/10.1111/j.1469-8986.2010.01087.x

Einhäuser, W., da Silva, L. F. O., & Bendixen, A. (2020). Intraindividual Consistency Between Auditory and Visual Multistability. Perception, 49(2), 119–138. https://doi.org/10.1177/0301006619896282

Einhäuser, W., Stout, J., Koch, C., & Carter, O. (2008). Pupil dilation reflects perceptual selection and predicts subsequent stability in perceptual rivalry. Proceedings of the National Academy of Sciences, 105(5), 1704–1709. https://doi.org/10.1073/pnas.0707727105

Feil, D., & Atmanspacher, H. (2010). Acategorial states in a representational theory of mental processes. J Consciousness Stud, 17(5 – 6), 72–101.

Fischl, B., Sereno, M. I., Tootell, R. B. H., & Dale, A. M. (1999). High-resolution intersubject averaging and a coordinate system for the cortical surface. Human Brain Mapping, 8(4), 272–284. https://doi.org/10.1002/(SICI)1097-0193(1999)8:4<272::AID-HBM10>3.0.CO;2-4

Foerster, F. R., Weibel, S., Poncelet, P., Dufour, A., Delevoye-Turrell, Y. N., Capobianco, A., Ott, L., & Giersch, A. (2021). Volatility of subliminal haptic feedback alters the feeling of control in schizophrenia. Journal of Abnormal Psychology, 130(7), 775–784. https://doi.org/10.1037/abn0000703

Friston, K., Zeidman, P., & Litvak, V. (2015). Empirical Bayes for DCM: A Group Inversion Scheme. Frontiers in Systems Neuroscience, 9. https://doi.org/10.3389/fnsys.2015.00164

Fuchs, M., Kastner, J., Wagner, M., Hawes, S., & Ebersole, J. S. (2002). A standardized boundary element method volume conductor model. Clinical Neurophysiology, 113(5), 702–712. https://doi.org/10.1016/S1388-2457(02)00030-5

Giles, N., Lau, H., & Odegaard, B. (2016). What Type of Awareness Does Binocular Rivalry Assess? Trends in Cognitive Sciences, 20(10), 719–720. https://doi.org/10.1016/j.tics.2016.08.010

Gramfort, A., Luessi, M., Larson, E., Engemann, D. A., Strohmeier, D., Brodbeck, C., Goj, R., Jas, M., Brooks, T., Parkkonen, L., & Hämäläinen, M. S. (2013). MEG and EEG data analysis with MNE-Python. Frontiers in Neuroscience, 7. https://doi.org/10.3389/fnins.2013.00267

Hecker, L., Tebartz van Elst, L., & Kornmeier, J. (2023). Source Localization Using Recursively Applied and Projected MUSIC with Flexible Extent Estimation [Preprint]. Neuroscience. https://doi.org/10.1101/2023.01.20.524892

Hecker, L., Wilson, M., Tebartz van Elst, L., & Kornmeier, J. (2022). Altered EEG variability on different time scales in participants with autism spectrum disorder: An exploratory study. Scientific Reports, 12(1), 13068. https://doi.org/10.1038/s41598-022-17304-x

Hense, M., Badde, S., & Röder, B. (2019). Tactile motion biases visual motion perception in binocular rivalry. Attention, Perception, & Psychophysics, 81(5), 1715–1724. https://doi.org/10.3758/s13414-019-01692-w

Huang, Q., Pereira, A. C., Velthuis, H., Wong, N. M. L., Ellis, C. L., Ponteduro, F. M., Dimitrov, M., Kowalewski, L., Lythgoe, D. J., Rotaru, D., Edden, R. A. E., Leonard, A., Ivin, G., Ahmad, J., Pretzsch, C. M., Daly, E., Murphy, D. G. M., & McAlonan, G. M. (2022). GABA B receptor modulation of visual sensory processing in adults with and without autism spectrum disorder. Science Translational Medicine, 14(626), eabg7859. https://doi.org/10.1126/scitranslmed.abg7859

Hupe, J. M., Lamirel, C., & Lorenceau, J. (2009). Pupil dynamics during bistable motion perception. Journal of Vision, 9(7), 10–10. https://doi.org/10.1167/9.7.10

Intaite, M., Koivisto, M., & Revonsuo, A. (2013). Perceptual reversals of Necker stimuli during intermittent presentation with limited attentional resources. Psychophysiology, 50(1), 82–96. https://doi.org/10.1111/j.1469-8986.2012.01486.x

Intaite, M., Koivisto, M., Ruksenas, O., & Revonsuo, A. (2010). Reversal negativity and bistable stimuli: Attention, awareness, or something else? Brain Cogn, 74(1), 24–34. https://doi.org/10.1016/j.bandc.2010.06.002

James, W. (1890). Principles of Psychology I.

Joos, E., Giersch, A., Hecker, L., Schipp, J., Heinrich, S. P., Tebartz van Elst, L., & Kornmeier, J. (2020). Large EEG amplitude effects are highly similar across Necker cube, smiley, and abstract stimuli. PLOS ONE, 15(5), e0232928. https://doi.org/10.1371/journal.pone.0232928

Kloosterman, N. A., Meindertsma, T., van Loon, A. M., Lamme, V. A. F., Bonneh, Y. S., & Donner, T. H. (2015). Pupil size tracks perceptual content and surprise. European Journal of Neuroscience, 41(8), 1068–1078. https://doi.org/10.1111/ejn.12859

Knapen, T., Brascamp, J., Pearson, J., van Ee, R., & Blake, R. (2011). The Role of Frontal and Parietal Brain Areas in Bistable Perception. Journal of Neuroscience, 31(28), 10293–10301. https://doi.org/10.1523/JNEUROSCI.1727-11.2011

Kornmeier, J., & Bach, M. (2004). Early neural activity in Necker-cube reversal: Evidence for low-level processing of a gestalt phenomenon. Psychophysiology, 41, 1–8. https://doi.org/doi: http://dx.doi.org/10.1016/j.visres.2004.10.006

Kornmeier, J., & Bach, M. (2006). Bistable perception—Along the processing chain from ambiguous visual input to a stable percept. Int J Psychophysiol, 62(2), 345–349.

Kornmeier, J., & Bach, M. (2012). Ambiguous figures – what happens in the brain when perception changes but not the stimulus. Frontiers in Human Neuroscience, 6(51), 1–23. https://doi.org/10.3389/fnhum.2012.00051

Kornmeier, J., & Bach, M. (2014). EEG correlates of perceptual reversals in Boring’s ambiguous old/young woman stimulus. Perception, 0(0), 0–0. https://doi.org/10.1068/p7741

Kornmeier, J., Ehm, W., Bigalke, H., & Bach, M. (2007). Discontinuous presentation of ambiguous figures: How interstimulus-interval durations affect reversal dynamics and ERPs. Psychophysiology, 44(4), 552–560.

Kornmeier, J., Friedel, E., Wittmann, M., & Atmanspacher, H. (2017). EEG correlates of cognitive time scales in the Necker-Zeno model for bistable perception. Consciousness and Cognition, 53, 136–150. https://doi.org/10.1016/j.concog.2017.04.011

Kornmeier, J., Friedel, Evelyn., Hecker, L., Schmidt, S., & Wittmann, M. (2019). What happens in the brain of meditators when perception changes but not the stimulus? PLOS ONE, 14(10), e0223843. https://doi.org/10.1371/journal.pone.0223843

Kornmeier, J., Hein, C. M., & Bach, M. (2009). Multistable perception: When bottom-up and top-down coincide. Brain and Cognition, 69(1), 138–147. https://doi.org/10.1016/j.bandc.2008.06.005

Kornmeier, J., Heinrich, S. P., Atmanspacher, H., & Bach, M. (2001). The reversing “Necker Wall” – a new paradigm with reversal entrainment reveals an early EEG correlate. ARVO 2001 Annual Meeting, 42, 409.

Kornmeier, J., Pfaffle, M., & Bach, M. (2011). Necker cube: Stimulus-related (low-level) and percept-related (high-level) EEG signatures early in occipital cortex. J Vis, 11(9), 1–11. https://doi.org/10.1167/11.9.12

Kornmeier, J., Wörner, R., Riedel, A., Bach, M., & Tebartz van Elst, L. (2014). A Different View on the Checkerboard? Alterations in Early and Late Visually Evoked EEG Potentials in Asperger Observers. PLoS ONE, 9(3), e90993. https://doi.org/10.1371/journal.pone.0090993

Kornmeier, J., Wörner, R., Riedel, A., & Tebartz van Elst, L. (2017). A different view on the Necker cube—Differences in multistable perception dynamics between Asperger and non-Asperger observers. PLOS ONE, 12(12), e0189197. https://doi.org/10.1371/journal.pone.0189197

Koshiyama, D., Kirihara, K., Tada, M., Nagai, T., Fujioka, M., Ichikawa, E., Ohta, K., Tani, M., Tsuchiya, M., Kanehara, A., Morita, K., Sawada, K., Matsuoka, J., Satomura, Y., Koike, S., Suga, M., Araki, T., & Kasai, K. (2018). Electrophysiological evidence for abnormal glutamate-GABA association following psychosis onset. Translational Psychiatry, 8(1), 211. https://doi.org/10.1038/s41398-018-0261-0

Krüger, J. (2022). Inattentive Perception, Time, and the Incomprehensibility of Consciousness. Frontiers in Psychology, 12, 804652. https://doi.org/10.3389/fpsyg.2021.804652

Lehmann, D., & Skrandies, W. (1980). Reference-free identification of components of checkerboard-evoked multichannel potential fields. Electroencephalography and Clinical Neurophysiology, 48(6), 609–621. https://doi.org/10.1016/0013-4694(80)90419-8

Leopold, D. A., & Logothetis, N. K. (1999). Multistable phenomena: Changing views in perception. aTrends Cogn Sci, 3(7), 254–264. https://doi.org/10.1016/S1364-6613(99)01332-7

Leopold, D. A., Wilke, M., Maier, A., & Logothetis, N. K. (2002). Stable perception of visually ambiguous patterns. Nat Neurosci, 5(6), 605–609.

Liaci, E., Bach, M., Tebartz van Elst, L., Heinrich, S. P., & Kornmeier, J. (2016). Ambiguity in Tactile Apparent Motion Perception. PLOS ONE, 11(5), e0152736. https://doi.org/10.1371/journal.pone.0152736

Long, G. M., & Toppino, T. C. (2004). Enduring interest in perceptual ambiguity: Alternating views of reversible figures. Psychological Bulletin, 130(5), 748–768. http://dx.doi.org/doi:10.1037/0033-2909.130.5.748

Lumer, E. D., Friston, K. J., & Rees, G. (1998). Neural correlates of perceptual rivalry in the human brain. Science, 280(5371), 1930–1934.

Maier, A., Wilke, M., Logothetis, N. K., & Leopold, D. A. (2003). Perception of temporally interleaved ambiguous patterns. Curr Biol, 13(13), 1076–1085.

Marques-Carneiro, J. E., Krieg, J., Duval, C. Z., Schwitzer, T., & Giersch, A. (2021). Paradoxical Sensitivity to Sub-threshold Asynchronies in Schizophrenia: A Behavioral and EEG Approach. Schizophrenia Bulletin Open, 2(1), sgab011. https://doi.org/10.1093/schizbullopen/sgab011

Nakatani, H., & van Leeuwen, C. (2013). Antecedent occipital alpha band activity predicts the impact of oculomotor events in perceptual switching. Frontiers in Systems Neuroscience, 7. https://doi.org/10.3389/fnsys.2013.00019

Necker, L. A. (1832). Observations on some remarkable optical phaenomena seen in Switzerland; and on an optical phaenomenon which occurs on viewing a figure of a crystal or geometrical solid. The London and Edinburgh Philosophical Magazine and Journal of Science, 1(5), 329–337. https://doi.org/10.1080/14786443208647909

Notredame, C.-E., Pins, D., Deneve, S., & Jardri, R. (2014). What visual illusions teach us about schizophrenia. Frontiers in Integrative Neuroscience, 8. https://doi.org/10.3389/fnint.2014.00063

Nunez, P. L., & Srinivasan, R. (2006). Electric Fields of the Brain. Oxford University Press. https://doi.org/10.1093/acprof:oso/9780195050387.001.0001

O’Donnell, B. F., Hendler, T., & Squires, N. K. (1988). Visual evoked potentials to illusory reversals of the Necker cube. Psychophysiology, 25(2), 137–143.

Orbach, J., Ehrlich, D., & Heath, H. (1963). Reversibility of the Necker cube: I. An examination of the concept of “satiation of orientation.舡 Percept Mot Skills, 17, 439–458.

Orbach, J., Zucker, E., & Olson, R. (1966). Reversibility of the Necker cube: VII.: Reversal rate as a function of figure-on and figure-off durations. Percept Mot Skills, 22, 615–618.

O’Shea, R. P., Kornmeier, J., & Roeber, U. (2013). Predicting visual consciousness electrophysiologically from intermittent binocular rivalry. PLoS ONE, 8(10), e76134. https://doi.org/10.1371/journal.pone.0076134

Pascual-Marqui, R. D. (1999). Review of Methods for Solving the EEG Inverse Problem. 1(1).

Pastukhov, A. (2016). Perception and the strongest sensory memory trace of multi-stable displays both form shortly after the stimulus onset. Attention, Perception, & Psychophysics, 78(2), 674–684. https://doi.org/10.3758/s13414-015-1004-4

Pastukhov, A., & Braun, J. (2007). Perceptual reversals need no prompting by attention. J Vision, 7(10)(5), 1–17.

Pastukhov, A., & Braun, J. (2008). A short-term memory of multi-stable perception. J Vis, 8(13), 7 1–14. https://doi.org/10.1167/8.13.7

Pastukhov, A., & Braun, J. (2013). Structure-from-motion: Dissociating perception, neural persistence, and sensory memory of illusory depth and illusory rotation. Attention, Perception, & Psychophysics, 75(2), 322–340. https://doi.org/10.3758/s13414-012-0390-0

Pastukhov, A., & Klanke, J.-N. (2016). Exogenously triggered perceptual switches in multistable structure-from-motion occur in the absence of visual awareness. Journal of Vision, 16(3), 14. https://doi.org/10.1167/16.3.14

Pearson, J., & Brascamp, J. (2008). Sensory memory for ambiguous vision. Trends Cogn Sci, 12(9), 334–341. https://doi.org/10.1016/j.tics.2008.05.006

Pitts, M. A., & Britz, J. (2011). Insights from intermittent binocular rivalry and EEG. Front Hum Neurosci, 5, 107. https://doi.org/10.3389/fnhum.2011.00107

Pitts, M. A., Gavin, W. J., & Nerger, J. L. (2008). Early top-down influences on bistable perception revealed by event-related potentials. Brain Cogn, 67(1), 11–24.

Pitts, M. A., Martínez, A., & Hillyard, S. A. (2010). When and where is binocular rivalry resolved in the visual cortex? Journal of Vision, 10(14), 25–25. https://doi.org/10.1167/10.14.25

Pitts, M. A., Nerger, J. L., & Davis, T. J. R. (2007). Electrophysiological correlates of perceptual reversals for three different types of multistable images. J Vision, 7(1), 1–14.

Polgári, P., Causin, J.-B., Weiner, L., Bertschy, G., & Giersch, A. (2020). Novel method to measure temporal windows based on eye movements during viewing of the Necker cube. PLOS ONE, 15(1), e0227506. https://doi.org/10.1371/journal.pone.0227506

Pressnitzer, D., & Hupe, J. M. (2006). Temporal dynamics of auditory and visual bistability reveal common principles of perceptual organization. Curr Biol, 16(13), 1351–1357. https://doi.org/10.1016/j.cub.2006.05.054

Robertson, C. E., Ratai, E.-M., & Kanwisher, N. (2016). Reduced GABAergic Action in the Autistic Brain. Current Biology, 26(1), 80–85. https://doi.org/10.1016/j.cub.2015.11.019

Sandberg, K., Barnes, G. R., Bahrami, B., Kanai, R., Overgaard, M., & Rees, G. (2014). Distinct MEG correlates of conscious experience, perceptual reversals and stabilization during binocular rivalry. NeuroImage, 100, 161–175. https://doi.org/10.1016/j.neuroimage.2014.06.023

Schirrmeister, R. T., Springenberg, J. T., Fiederer, L. D. J., Glasstetter, M., Eggensperger, K., Tangermann, M., Hutter, F., Burgard, W., & Ball, T. (2017). Deep learning with convolutional neural networks for EEG decoding and visualization: Convolutional Neural Networks in EEG Analysis. Human Brain Mapping, 38(11), 5391–5420. https://doi.org/10.1002/hbm.23730

Schmack, K., Schnack, A., Priller, J., & Sterzer, P. (2015). Perceptual instability in schizophrenia: Probing predictive coding accounts of delusions with ambiguous stimuli. Schizophrenia Research: Cognition, 2(2), 72–77. https://doi.org/10.1016/j.scog.2015.03.005

Sergent, C., Faugeras, F., Rohaut, B., Perrin, F., Valente, M., Tallon-Baudry, C., Cohen, L., & Naccache, L. (2017). Multidimensional cognitive evaluation of patients with disorders of consciousness using EEG: A proof of concept study. NeuroImage: Clinical, 13, 455–469. https://doi.org/10.1016/j.nicl.2016.12.004

Sforza, C., Rango, M., Galante, D., Bresolin, N., & Ferrario, V. F. (2008). Spontaneous blinking in healthy persons: An optoelectronic study of eyelid motion. Ophthalmic and Physiological Optics, 28(4), 345–353. https://doi.org/10.1111/j.1475-1313.2008.00577.x

Sterzer, P., & Kleinschmidt, A. (2007). A neural basis for inference in perceptual ambiguity. Proc Natl Acad Sci U S A, 104(1), 323–328.

Strüber, D., & Herrmann, C. S. (2002). MEG alpha activity decrease reflects destabilization of multistable percepts. Cogn Brain Res, 14(3), 370–382.

Strüber, D., & Stadler, M. (1999). Differences in top-down influences on the reversal rate of different categories of reversible figures. Perception, 28, 1185–1196.

Tong, F., Nakayama, K., Vaughan, J. T., & Kanwisher, N. (1998). Binocular rivalry and visual awareness in human extrastriate cortex. Neuron, 21(4), 753–759.

van Ee, R., van Dam, L. C., & Brouwer, G. J. (2005). Voluntary control and the dynamics of perceptual bi-stability. Vision Res, 45(1), 41–55.

VanRullen, R., & Thorpe, S. J. (2002). Surfing a spike wave down the ventral stream. Vision Res, 42(23), 2593–2615.

Watanabe, T. (2021). Causal roles of prefrontal cortex during spontaneous perceptual switching are determined by brain state dynamics. ELife, 10, e69079. https://doi.org/10.7554/eLife.69079

Weilnhammer, V., Fritsch, M., Chikermane, M., Eckert, A.-L., Kanthak, K., Stuke, H., Kaminski, J., & Sterzer, P. (2021). An active role of inferior frontal cortex in conscious experience. Current Biology, 31(13), 2868–2880.e8. https://doi.org/10.1016/j.cub.2021.04.043

Weilnhammer, V., Stuke, H., Hesselmann, G., Sterzer, P., & Schmack, K. (2017). A predictive coding account of bistable perception—A model-based fMRI study. PLOS Computational Biology, 13(5), e1005536. https://doi.org/10.1371/journal.pcbi.1005536

Wexler, M. (2018). Multidimensional internal dynamics underlying the perception of motion. Journal of Vision, 18(5), 7. https://doi.org/10.1167/18.5.7

Yokota, Y., Minami, T., Naruse, Y., & Nakauchi, S. (2014). Neural processes in pseudo perceptual rivalry: An ERP and time–frequency approach. Neuroscience, 271, 35–44. https://doi.org/10.1016/j.neuroscience.2014.04.015

Zaretskaya, N., Thielscher, A., Logothetis, N. K., & Bartels, A. (2010). Disrupting parietal function prolongs dominance durations in binocular rivalry. Curr Biol, 20(23), 2106–2111. https://doi.org/10.1016/j.cub.2010.10.046

